# Serotype specific sugars impact structure but not functions of the trimeric autotransporter adhesin EmaA of *Aggregatibacter actinomycetemcomitans*

**DOI:** 10.1101/2022.06.07.495235

**Authors:** Gaoyan G. Tang-Siegel, Michael Radermacher, Keith P. Mintz, Teresa Ruiz

## Abstract

The human oral pathobiont *Aggregatibacter actinomycetemcomitans* expresses multiple virulence factors including the trimeric, extracellular matrix protein adhesin A (EmaA). The posttranslational modification of EmaA is proposed to be dependent on the sugars and enzymes associated with O-polysaccharide (O-PS) synthesis of the lipopolysaccharide (LPS). This modification is important for the structure and function of this adhesin. To determine if the composition of the sugars alters structure and/or function, the prototypic 202 kDa protein was expressed in a non-serotype b, *emaA* mutant strain. The transformed strain displayed EmaA adhesins similar in appearance to the prototypic adhesin as observed by 2D electron microscopy of whole-mount negatively stained bacterial preparations. Biochemical analysis indicated that the protein monomers were post-translationally modified. 3D electron tomographic reconstruction and structure analyses of the functional domain revealed three well-defined subdomains (SI, SII and SIII) with a linker region between SII and SIII. Structural changes were observed in all three subdomains and the linker region of the adhesins synthesized compared with the known structure. These changes however did not affect the ability of the strain to bind collagen or form biofilms. The data suggest that changes in the composition of the glycan moiety alter the 3D structure of the molecule without negatively affecting the function(s) associated with this adhesin.

**IMPORTANCE:** The human oral pathogen *A. actinomycetemcomitans* is a causative agent of periodontal and several systemic diseases. EmaA is a trimeric autotransporter protein adhesin important for the colonization of this pathobiont *in vivo*. This adhesin is modified with sugars associated with the O-polysaccharide (O-PS) and the modification is mediated using the same enzymes involved in lipopolysaccharide (LPS) biosynthesis. The interaction with collagen is not mediated by the specific binding between the glycans and collagen but is attributed to changes in the final quaternary structure necessary to maintain an active adhesin. In this study, we have determined that the composition of the sugars utilized in the post-translational modification of this adhesin is exchangeable without compromising functional activities.

## INTRODUCTION

Bacterial adhesion is mediated by proteinaceous surface structures including fimbrial and non-fimbrial adhesins. The <u>e</u>xtracellular <u>m</u>atrix protein <u>a</u>dhesin <u>A</u> (EmaA) expressed by the human oral pathogen *Aggregatibacter actinomycetemcomitans* is a non-fimbrial adhesin promoting colonization (1–3). EmaA is classified as a trimeric autotransporter protein (type V_c_ secretion system) and belongs to a family of proteins, which is represented by YadA of *Yersinia enterocolitica* (1). Three monomers of EmaA form antenna-like projections, extending up to 150 nm from the bacterial outer membrane as visualized by transmission electron microscopy (TEM) (4, 5).

The functional domain of the adhesin is associated with the ellipsoidal end of the projections which corresponds to the first 760 amino acids of the protein monomer after cleavage of the 56 amino acid signal peptide (5–8). The remainder of the sequence forms the stalk, including the membrane pore-forming domain (4, 5). The distal end of the structure is further subdivided into three subdomains (SI, containing amino acids 57-225, SII containing amino acids 226-433, and SIII containing amino acids 448-627) with a linker region between SII and SIII containing 14 amino acids (6, 8). Deletion of the whole functional domain or specific subdomains negatively affects the collagen binding activity of this adhesin (5, 8). The functional domain is also involved in cell-cell interactions in microcolony development during biofilm formation (3).

The *emaA* gene is ubiquitous in *A. actinomycetemcomitans*, however, heterogeneity exists in the protein sequence that correlates with the serotype of the strain (9). Serotypes b and c strains express the prototypic (202 kDa, b-EmaA) protein, whereas serotypes a and d strains encode a 173 kDa (a-EmaA) homolog. The a-EmaA isomer shares 75% amino acid sequence similarity in the functional domain and 95% sequence similarity within the stalk and pore-forming sequence. The reduction in the size of the monomer is due to a 279 amino acid deletion in the region between the head and stalk domains of the prototypic sequence (9). Both protein species form structures that bind to collagen and participate in biofilm formation (3, 9).

Biochemical and genetic studies support the hypothesis that the b-EmaA is glycosylated with O-polysaccharide (O-PS) sugars mediated by the Waal ligase of the lipopolysaccharide (LPS) pathway (10). This posttranslational modification is necessary for the collagen binding activity and protein stability but is not essential for biofilm formation (3, 10–12). The glycan is not directly involved in collagen binding but is required for the functional quaternary structure of the adhesin. However, the role of the specific glycan composition in the structural stability and function of the adhesin remains unknown.

To analyze the functional and structural role of the sugars modifying EmaA, we have modified the glycan moiety by transforming a serotype a strain (O-PS composed of D-talose disaccharide repeats), which does not express the endogenous a-EmaA protein, with a plasmid encoding the prototypic b-EmaA protein sequence under the control of the endogenous promoter. The transformed strain displayed antenna-like projections, similar to prototypic EmaA adhesins expressed in serotype b strains, as visualized by TEM, and the EmaA monomers displayed characteristics of a post-translationally modified protein. Furthermore, electron tomography and subvolume averaging were utilized to determine the substructure of the functional region of this prototypic b-EmaA expressed in the serotype a strain. The overall structure was found to be similar to the prototypic b-EmaA, but clear differences in the density of the subdomains and the linker region were observed. Despite the difference in EmaA structures, the transformed strain was active in binding to collagen and forming biofilms. The data suggest that the composition change of associated glycans does not influence the known functions of this adhesin.

## RESULTS

### Transmission electron microscopy images of the *A. actinomycetemcomitans* serotype a strain transformed with the prototypic b-EmaA

The serotype a strain transformed by electroporation with the plasmid expressing the full-length b-*emaA* gene (pKM11) under control of the endogenous promoter sequence was examined by 2D electron microscopy (2DEM) of whole-mount negatively stained bacterial preparations. Antenna-like projections were observed associated with the rugose membrane of the transformed cells. These surface projections extended well away from the bacterial surface and contained bends similar to those observed in the serotype b wild type strain expressing the chromosomal copy of the gene. The increased number of projections associated with the transformed strain ATCC29523/pKM11 (FIG 1A) is due to the over-expression of the gene associated with the plasmid. The overall structure is similar to the prototypic EmaA expressed in serotype b (FIG 1B).

**FIG 1.**
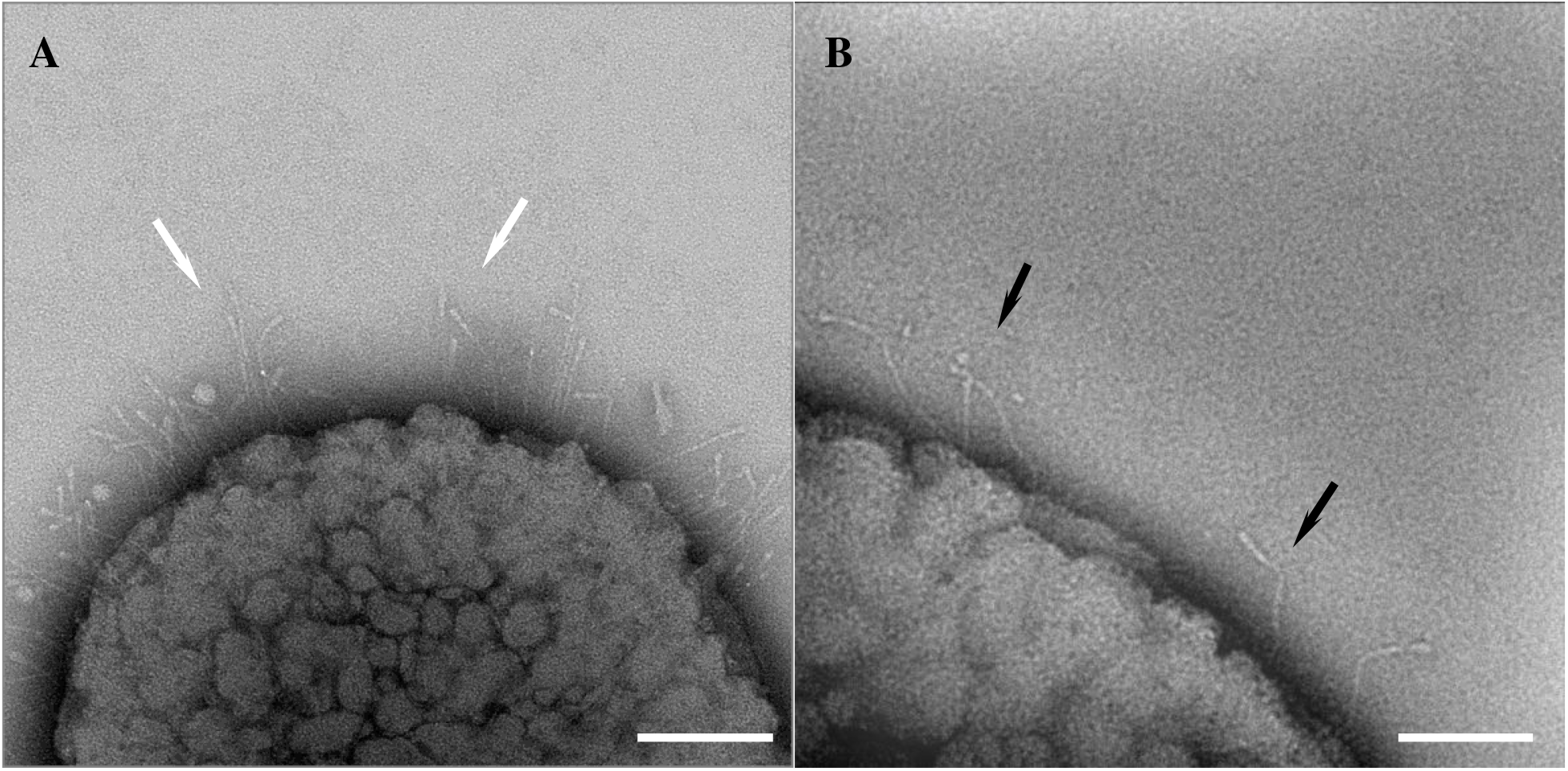
Transmission electron micrographs of b-EmaA expressed in different O-PS backgrounds. **A)** The EmaA structures (shown by white arrows) were expressed in a serotype a *emaA* mutant strain (ATCC29523) using the plasmid pKM11. **B**) The prototypic b-EmaA (shown by black arrows) expressed in the serotype b strain VT1169. Scale bar = 100 nm.

### Biochemical characterization of EmaA

The electrophoretic mobility of the EmaA monomers isolated from the membrane of the transformed serotype a strain was similar to the mobility of the monomers isolated from the serotype b wild type strain (FIG 2A). The monomers associated with both strains had a reduced mobility when compared with the monomers derived from the membrane of a serotype b strain deficient in O-PS synthesis (*rmlC^-^*) (FIG 2A). The immunoreactive material at the top of the wells represents aggregates of EmaA that do not enter the running gel, which is also consistent with the biochemical feature of this adhesin (10, 13). No immunoreactive material was observed in the membrane fraction of the serotype a bacteria transformed with the empty plasmid. In addition, a comparison of the EmaA monomers isolated from the cytosol with those associated with the membrane demonstrated a difference in the apparent electrophoretic mobilities (FIG 2B). This difference is indicative of a change in the mass of the protein. Taken together, the data suggest that the monomers isolated from the serotype a strain transformed with b-*emaA* resemble the characteristics of the wild type monomers isolated from serotype b strain.

**FIG 2.**
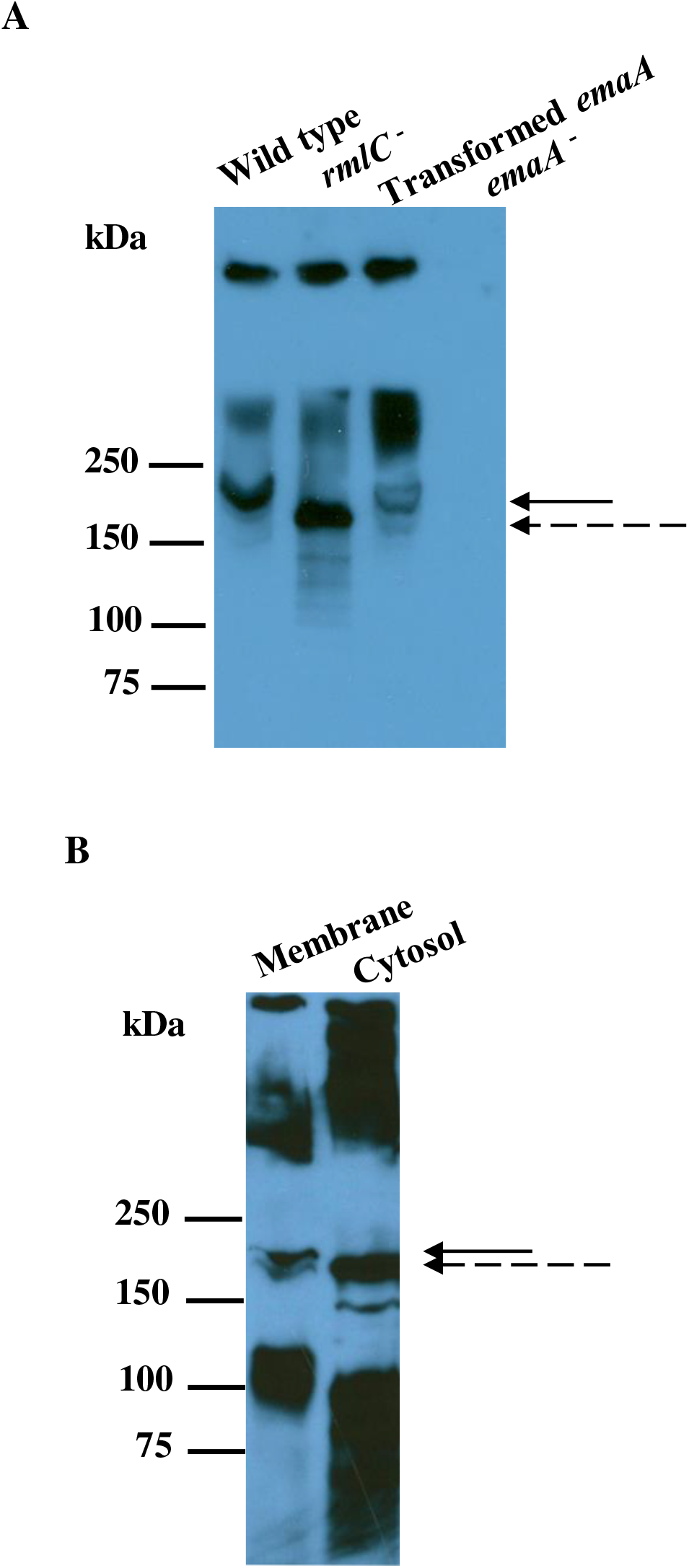
Biochemical characterization of EmaA expressed in different O-PS backgrounds. **A)** Immunoblot of membrane EmaA monomers in different O-PS backgrounds. Wild type: VT1169 (serotype b); *rmlC*^-^: the isogenic *rmlC* mutant of VT1169; Transformed EmaA: a serotype a *emaA* mutant strain ATCC29523 transformed with plasmid expressing b-*emaA* (pKM11); *emaA*^-^: ATCC29523 transformed with the empty plasmid (pKM2). **B)** Immunoblot of cytosol and membrane localized EmaA of the transformed *emaA* strain (ATCC29523/pKM11). Solid arrows indicate electrophoretic mobility of the membrane and dashed arrows indicate electrophoretic mobility of monomers isolated from an O-PS mutant (*rmlC^-^*) (A) or from the cytosol (B).

### 3D structure of the EmaA functional domain

The structure of the EmaA functional domain was determined by electron tomography and subvolume averaging techniques of whole-mount negatively stained bacterial preparations of the transformed serotype a strain. 84 tomographic single-axis tilt series were collected with an angular range of at least ± 64° in 2° increments, the defocus was selected so that the first zero of the contrast transfer function (CTF) would lie at a resolution better than 15 Å. A total of 151 EmaA functional domains were selected from the tomograms (FIG 3A) and reconstructed from the corresponding subprojections using Radon transform algorithms (FIG 3B) (14). After an initial interactive alignment, using the prototypic EmaA as reference, 15% of the subvolumes were discarded due to the close proximity of either fiducials or proteinaceous contamination. The data was subjected to a first round of classification using principal component analysis (PPCA-EM subroutine implemented in EMIRA) (15, 16) followed by clustering to aid subsequent alignment steps within this dataset. A representative for each subclass was used as a new reference in an interactive multireference alignment step. Since the data exhibited conformational heterogeneity, a final round of classification was performed in the remaining 128 EmaA subvolumes and eight classes were obtained with different numbers (FIG 4).

**FIG 3.**
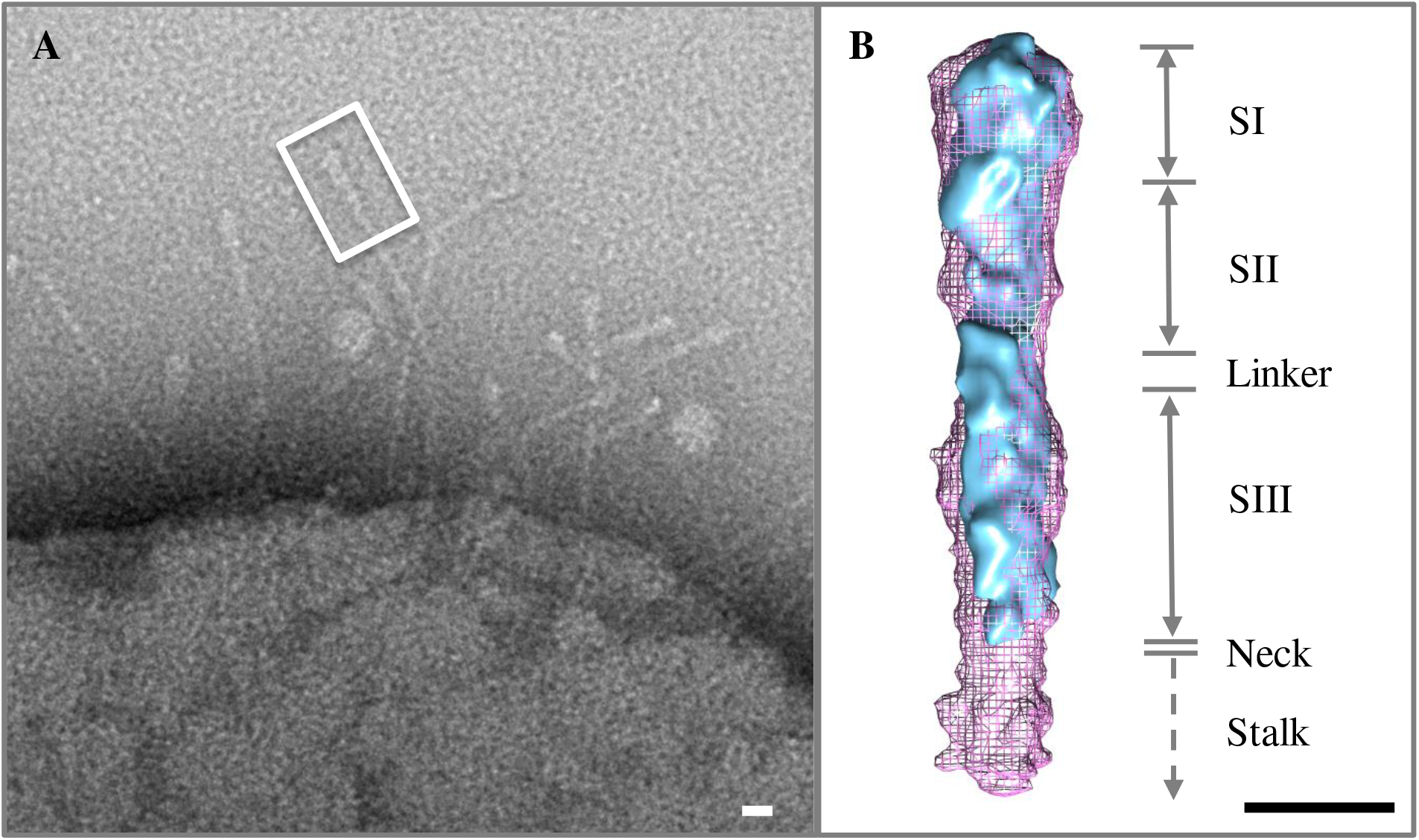
Functional N-terminus of b-EmaA expressed in the serotype a transformed strain *vs* the wild-type EmaA expressed in a serotype b strain. **A)** The antenna-like EmaA projections are visualized extending from the bacterial surface. The functional N-terminus is approximately 30 nm from the distal end of the EmaA appendage (shown in the white square). **B)** Structures of the functional N-terminus: the average wild type EmaA (pink mesh) *versus* one individual transformed EmaA (blue surface). The functional terminus includes three subdomains: SI (amino acids 57-225), SII (226-433) and SIII (448-627) and a more flexible linker region between SII and SIII. Scale bar = 5 nm.

**FIG 4.**
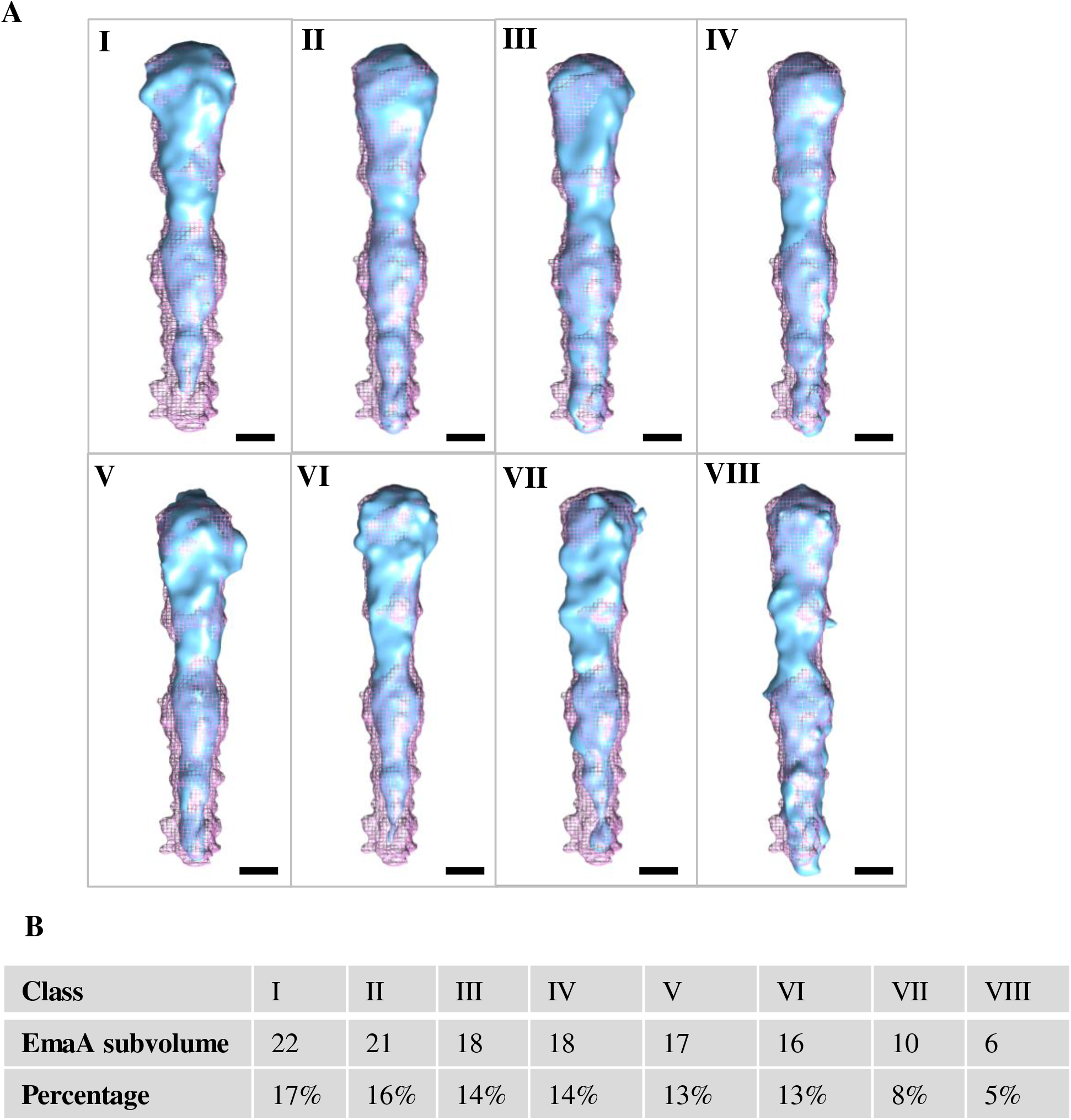
Surface representation of subvolume averages of the functional N-terminus of the b-EmaA expressed in the transformed serotype a strain. **A)** The average EmaA subvolume of each subclass, in a total of eight subclasses based on 128 EmaA subvolumes extracted from 84 reconstructed tomograms, is demonstrated in blue surface *versus* the wild type EmaA (from the serotype b strain) in pink mesh, **B)** Distribution of eight different subclasses in the b-EmaA expressed in the transformed serotype a strain. Scale bar = 3 nm.

The overall structure of the EmaA subvolumes (from the serotype a transformed strain with the serotypeb-emaA gene) calculated from the combined subprojection sets displayed a similar subdomain architecture as the prototypic b-EmaA average (FIG 4A and Movie1 in Supplementary Materials). However, in general, subdomain SII in all classes was significantly smaller than in the prototypic EmaA, this effect is more visible in classes I and II. Moreover, 73% of the EmaA population had an enlarged subdomain SI. This appears most evident in classes V and VI, which have concomitantly smaller density in subdomains SII and SIII. An additional feature present in a few of these classes (*e.g.* III, IV, VII and VIII) is a bend situated between the linker region and subdomain SIII, which rarely appears in the prototypic EmaA (5). An analysis of direct class averages and variances calculated from the reconstituted volumes after the last round of PPCA-EM revealed the largest variance to be located at the core of the structure within subdomains SI and SII (data not shown). Only class II and to a minor extent classes V and VIII show variances on the external surface of the adhesin as it would have been expected.

Volume segmentation was carried out to characterize the continuity of the density along the functional domain of the EmaA adhesins (17). Segmentation of the EmaA class average subvolumes in Chimera using three smoothing steps resulted in subdivisions of the SI-SIII region that contained two segments as for the prototypic EmaA either with or without 3-fold symmetrization (FIG 5 and Movie2 in Supplementary Materials). However, the results of the segmentation differed in the specific start and end of the segments. The first prototypic EmaA segment contained SI, SII and a third of the linker region (FIG 5, WT) while the b-EmaA expressed in the serotype a strain contained SI, SII and either half of the linker region for classes II, III and VIII or the complete linker region for the rest of the classes. Taken together, these data indicate that there are differences between the prototypic b-EmaA and the EmaA structures when expressed in different O-PS backgrounds.

**FIG 5.**
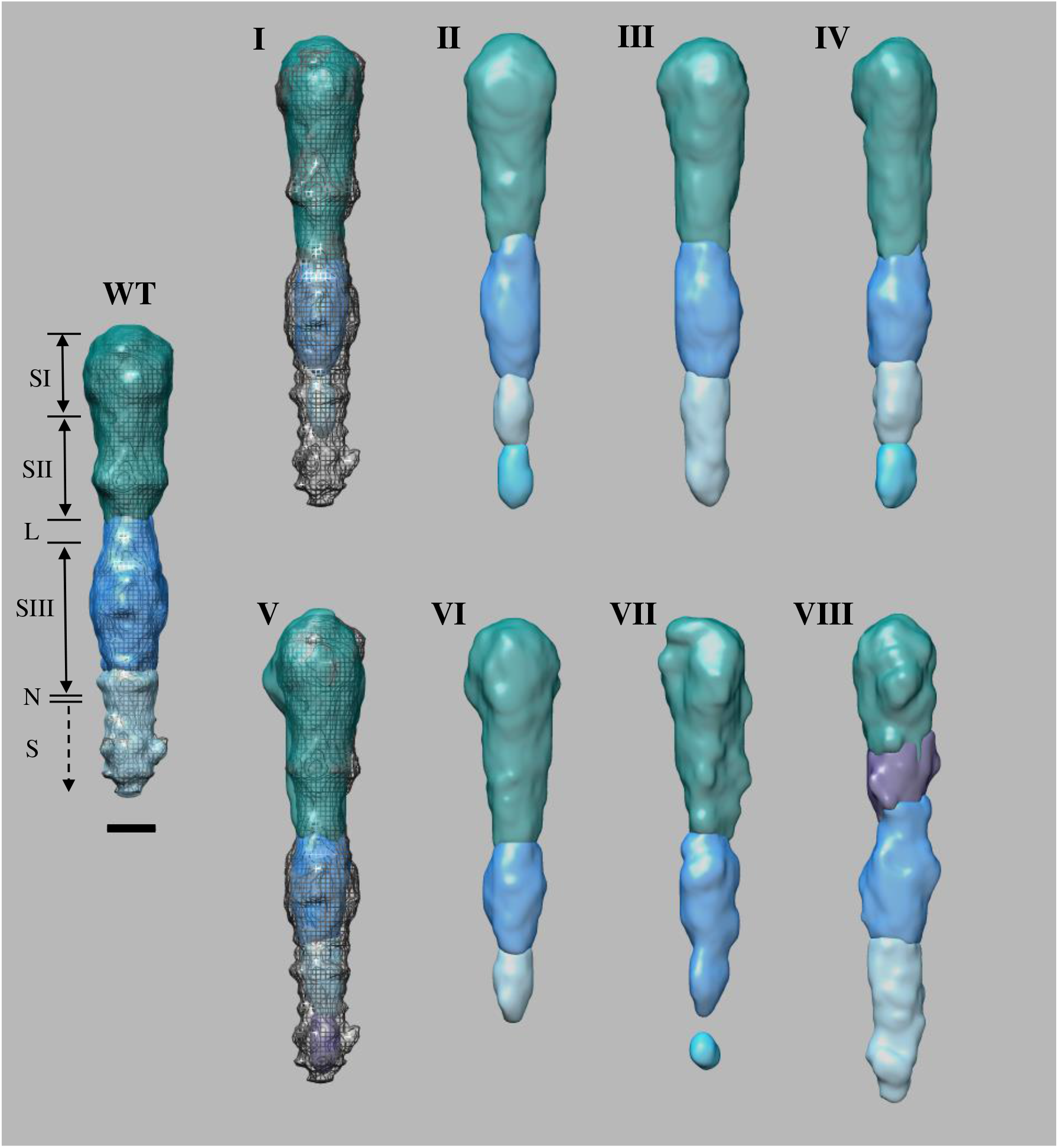
Segmentation of subvolume averages of the functional N-terminus of the b-EmaA expressed in the transformed serotype a strain. Segmentation of the subvolume average of each subclass of the b-EmaA, expressed in the transformed serotype a strain. Segmentation was carried out using three smoothing steps and resulted in subdivisions of the functional region of EmaA between SI and SIII. The wild type EmaA (serotype b) has been labelled WT and the subclasses are labelled I to VIII as in FIG 4. SI, SII, SIII represent three well-defined subdomains of the structure. L is the Linker region between SII and SIII; N is a Neck sequence and S is the Stalk domain. Scale bar = 3 nm.

### Functional activities of EmaA expressed in a non-cognate serotype strain

Changes in the structure of the functional domain may correlate with an alteration in the functions of the adhesin. The transformed serotype a strain expressing the b-*emaA* was observed to form a more robust biofilm compared to the strain transformed with the empty plasmid (FIG 6). The biofilm formed was 5- and 10-fold greater in mass at 48 and 72 hours respectively, suggesting that changes in structures did not impact the ability of the strain expressing EmaA to mediate biofilm formation. Collagen binding activity, however, is dependent on the posttranslational modification. In collagen binding assays, a 3-fold increase in collagen binding activity of the transformed strain was observed when compared with the parent mutant strain (FIG 6). These data demonstrate that b-EmaA modified with D-talose still retains collagen binding activity.

**FIG 6.**
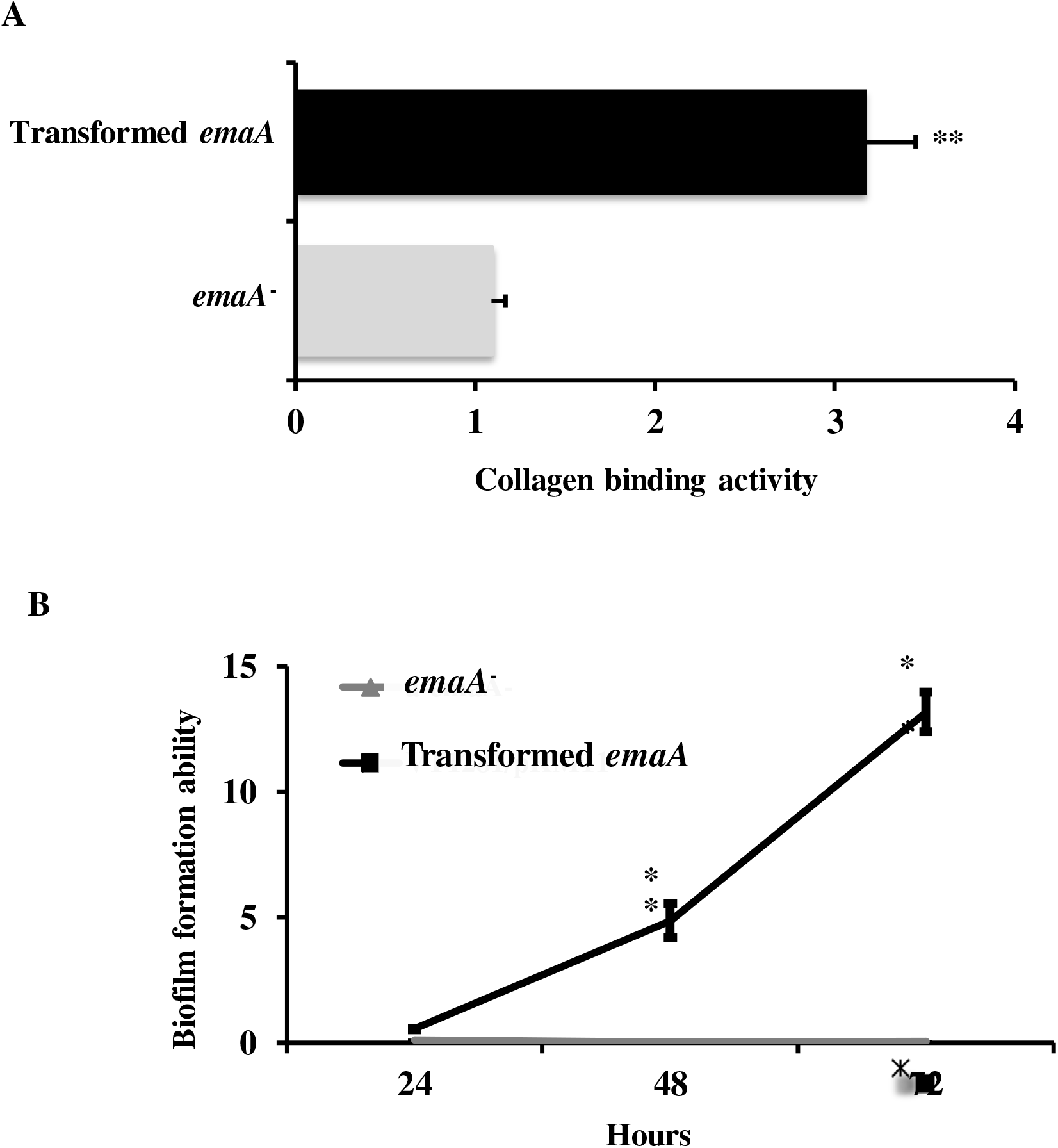
Functional analyses of b-EmaA expressed in the transformed serotype a strain. **A)** Collagen binding activity (**: *P* < 0.01). **B)** Biofilm formation. *emaA*^-^: serotype a *emaA* mutant strain transformed with an empty plasmid. Transformed *emaA*: serotype a strain transformed with a plasmid expressing b-*emaA* (*: *P* < 0.05).

## DISCUSSION

The multifunctional collagen adhesin EmaA is a recognized virulence factor contributing to the early colonization of the oral pathogen *A. actinomycetemcomitans* (1–3). EmaA is proposed to be glycosylated using a novel mechanism via the O-PS biosynthetic pathway (10). The posttranslational modification is required for collagen binding activity (13) and there is mounting evidence that the sugars are required for the formation of the quaternary structure of the adhesin (11, 12). To investigate the role of specific sugars in the folding of the protein, we have substituted the sugars of the serotype b (branched L-Rha, D-Fuc and D-GalNAc) with those expressed in serotype a strain (disaccharide units of D-Talose), which does not express EmaA on the surface due to the presence of an amber mutation in the 3’ end of the gene (9).

The 2D electron microscopy images of the serotype a *emaA* mutant strain transformed with a replicating plasmid containing the endogenous promoter with the intact b-*emaA* gene sequence validated the competency of the strain to express and form EmaA surface structures. These structures resembled those of the prototypic adhesin forming antenna-like appendages and contained multiple bends along the length of the structures, which are important for the flexibility of the adhesin (4–6, 18).

The electrophoretic mobility of the b-EmaA monomers expressed in the serotype a strain is similar to the prototypic b-EmaA expressed in the cognate serotype b background. The molecular mass of the protein expressed in the serotype a strain is also greater than the prototypic monomer expressed in a glycosylation deficient mutant strain (FIG 2) (10, 13). The glycosylation state of the protein is re-enforced by the difference in the electrophoretic mobility of the membrane associated form and the monomer found in the cytoplasm (FIG 2B), and it is consistent with the protein monomers isolated from these two fractions associated with the prototypic EmaA expressed in the serotype b background (13). Direct evidence for the glycosylation state of the protein is lacking due to difficulties in purifying membrane bound EmaA adhesins. This glycosylated trimeric adhesin tends to form aggregates trapped in the stacking SDS-PAGE gel, with only small portions of monomers running into the separating gel, which was demonstrated in the prototypic EmaA expressed by the wild type serotype b strain (10, 13), as well as here with the b-EmaA expressed in serotype a strains (FIG 2). Additional support stems from the structural data. Adhesins expressed on O-PS mutant strains seem to “hug” the cell surface, which might ensue from modifications of the electrostatic properties of both surfaces and exhibit a strong curvature along the whole length of the functional domain (13). The adhesins in this study extend well away from the bacterial surface (FIG 1A) and have relatively straight functional domains (FIGs. 3, 4 and 5). Thus, the biochemical and structural data presented in this study suggest that the prototypic protein expressed in this transformed serotype a strain is glycosylated.

The serotype a modified proteins also displayed the activities associated with functional EmaA adhesins. The transformed strain demonstrated greater collagen binding and biofilm formation as compared with the parent strain, which is a spontaneous *emaA* mutant. Collagen binding is dependent on the glycosylation state of the protein based on the decrease in binding activity of strains mutant for OPS metabolic, transport or ligase enzymes (10, 13). Since binding is dependent on the posttranslational modification, the increase in activity lends additional support to the nature of the modification and the role of this modification in the formation of an active adhesin. The adhesin is also important in biofilm biogenesis (3) and the functionality of the adhesin is further supported by the observation of increased biofilm biogenesis of the transformed strain (FIG 6B). Although biofilm formation is independent of the glycosylation state and the precise 3D structure (3, 6, 8), expression of the b-EmaA modified with different sugars does not affect the activity of this adhesin.

The collagen binding and biofilm formation abilities of the transformed strain in this study (FIG. 6) implies that it is glycosylation itself that is relevant for the proper function of EmaA, rather than the specific sugars bound to the protein. In *E. coli*, O-PS is synthesized as undecaprenyl-diphosphate (Und-PP)-linked O-antigen, transferred from the cytoplasm to the periplasmic space, and covalently ligated to the lipid A-core oligosaccharide using the WaaL ligase (19). Since EmaA utilizes the WaaL ligase for glycosylation, it indicates that the linkages between glycans and the protein involve at least O-linked glycosylation. The potential sites for O-linked glycosylation, Ser and Thr residues are widely distributed across the whole protein. Glycosylation is required for EmaA binding to collagen and stability of the protein (11–13). A comparison between glycosylation deficient and prototypic b-EmaA structures (6, 8, 11, 12, 17) reveals structural changes within the functional domain, including subdomains I, II and III and the linker region, as demonstrated in the serotype a modified protein (FIGs. 4 and 5). Therefore, taken together these data indicate that the functional domain of EmaA contains the glycosylation sites (FIG. 7).

**FIG 7.**
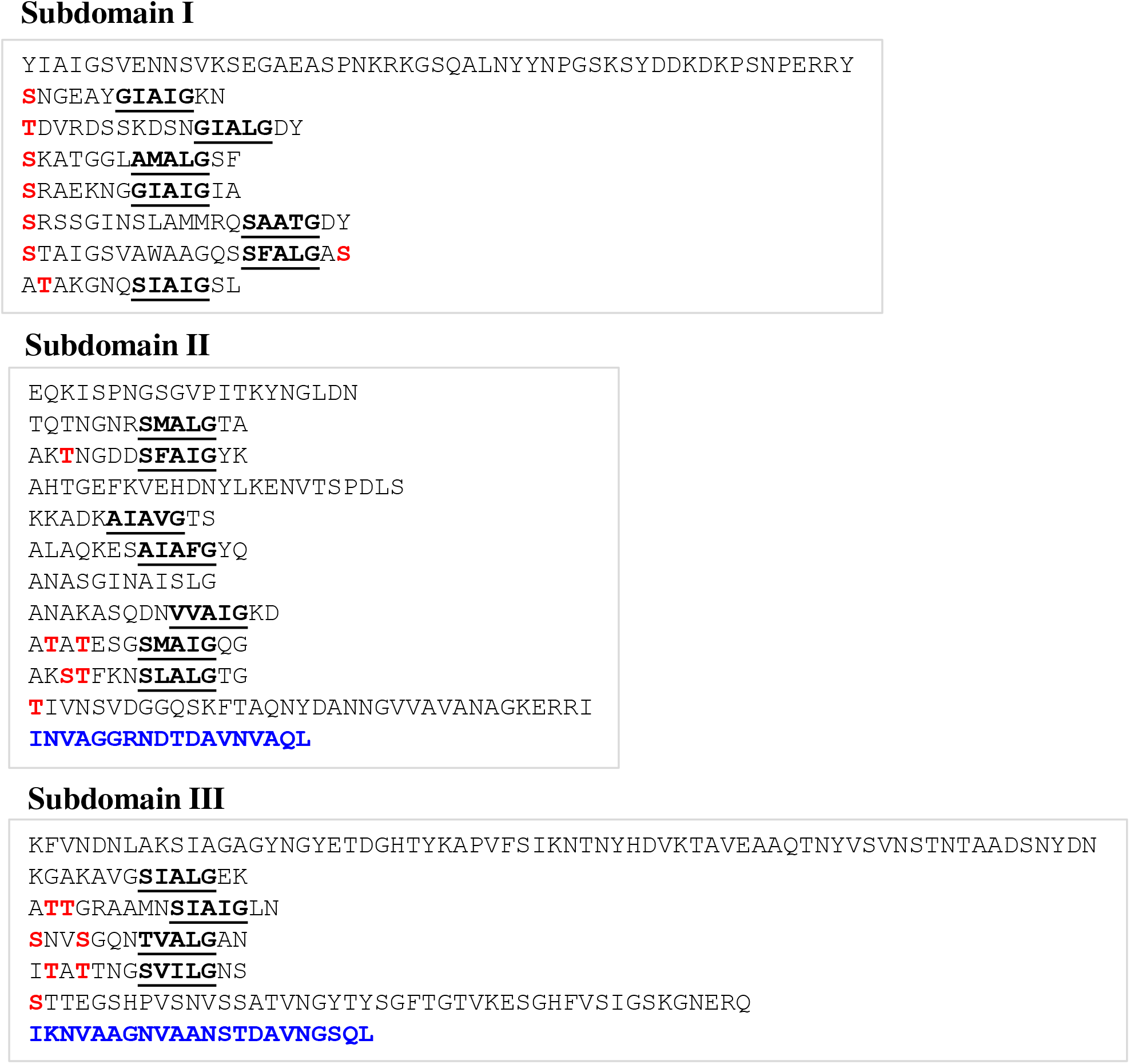
Predicted O-linked glycosylation sites of the EmaA functional domain. Sequence of the N-terminal 627 amino acids of EmaA: Subdomain I (amino acids 57-225), Subdomain II (226-433) and Subdomain III (448-627). Pentameric repeats are bold-underlined, (– **<u>SVAIG</u>**XXS-). Two neck sequences are highlighted in blue. The possible O-linked glycosylation sites (highlighted in red) are S(Ser) or T(Thr), two amino acids after the pentameric repeats.

The structural integrity of the EmaA adhesin has been shown to be critical for optimal function (6, 8, 11, 12). Deletion mutants of the protein sequence located in the functional domain (subdomains SI-SIII) decrease or abolish the collagen binding ability of the adhesin. Even the mutation of a single amino acid (G162S) located at the core of the structure resulted in a negative functional outcome and concomitantly in a structural change (6, 8). The bend in the adhesin between subdomains SII and SIII observed in the transformed strain in this study was reminiscent of the bend displayed by the adhesin bound to collagen (8, 18). Mutants defective in the O-PS biosynthetic pathway (*rmlC* and *waaL* mutant strains) exhibited overall a reduced density of the functional domain. Specifically, subdomain SI had a diameter comparable to subdomain SII (∼ 4.4 nm) instead of the prototypical SI diameter of 5 nm. In this study, we observed an enlarged subdomain SI concomitantly with weaker densities in subdomains SII and SIII. The larger volume occupied by subdomain SI might be the result of either an increased density or a more loosely packed structure resulting from the different sugars. In addition, the functional domain of O-PS mutants presented a strong curvature and multiple bends not commonly observed in the prototypical EmaA (11, 12). None of those bends were observed in the EmaA structures in the present study (FIG 4). The data from this study indicate that the correlation between structure and function of this adhesin is less straightforward than originally suggested and that a certain structural permissiveness is allowed for proper function.

The nature of the sugars post-translationally modifying EmaA affects the continuity of the density along the functional domain of the adhesins. Classification and segmentation of subvolumes calculated from 3D reconstructions of the prototypic EmaA shows that the SI and SII segment together independently of subdomain SIII (FIG 5, WT). Additionally, the top SI/SII segment contains a third of the linker region while the bottom SIII segment contains the remaining two thirds of the linker region. In most of the classes of this study (except for Class VIII) we observe that the whole linker region is part of the top SI/SII segment with the bottom segment containing mainly SIII. This indicates that the different sugars modifying the EmaA protein play a role in the interconnectivity and intraconnectivity of the subdomains, which are relevant for the functional activity of the adhesin.

In summary, we have determined that changes in the composition of the glycan moiety associated with the protein does not impact the function of this multifunctional adhesin. However, the data suggests that the overall 3D structure of the adhesin is dependent on the composition of the glycan moiety, without concomitantly negatively affecting the function(s) associated with EmaA.

## MATERIALS AND METHODS

### Strains and plasmids

All bacterial strains used in this study are listed in Table 1. *A. actinomycetemcomitans* strains were grown statically in 3% trypticase soy broth, 0.6% yeast extract (TSBYE), with or without 1.5% agar (Becton Dickinson and Company) and the required antibiotics in a humidified 37 °C incubator with 5% carbon dioxide. All mutant strains in this study retained growth characteristics similar to the wild type strain. *Escherichia coli* was grown in broth containing 1% BactoTryptone, 0.5% yeast extract and 1% sodium chloride (Lysogenic broth, LB) with appropriate antibiotics at 37 °C under aerobic conditions with agitation. All sequencing was performed at the University of Vermont Cancer Center DNA Analysis Facility. The shuttle plasmid pKM2 was used for transformation and expression of EmaA (20).

**TABLE 1.**
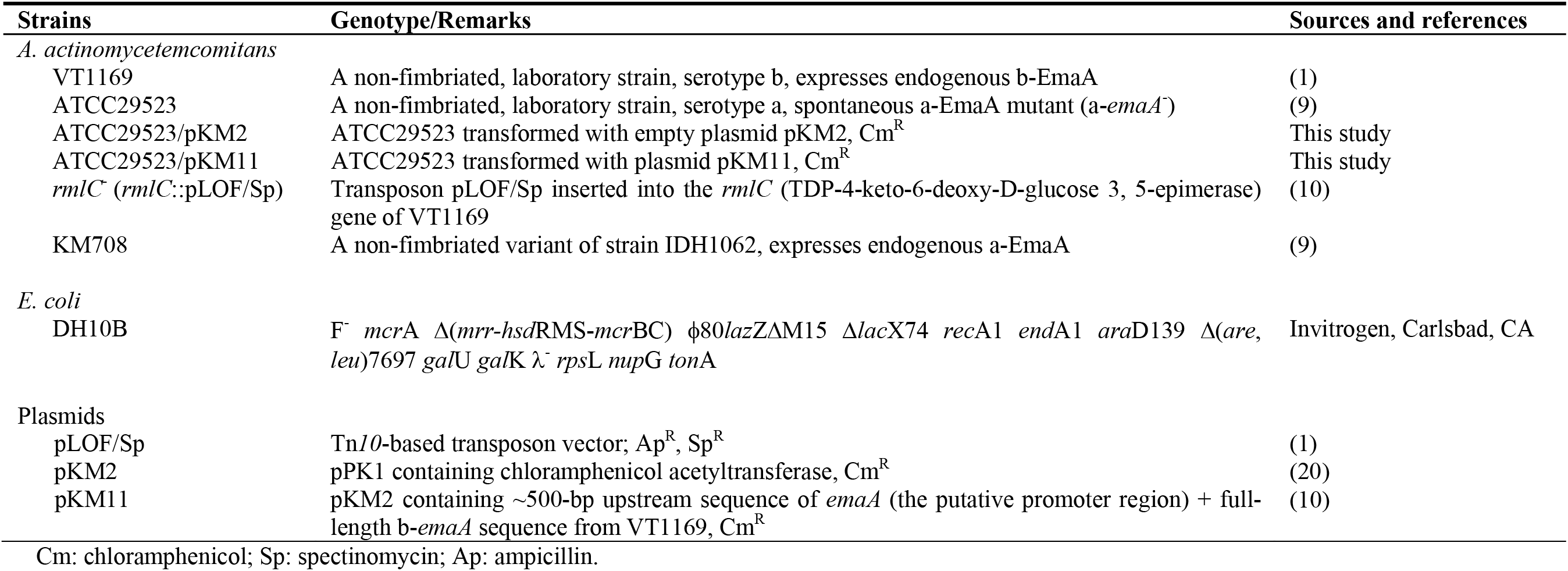
Strains and plasmids.

### Fractionation of membrane and cytosol proteins

Membrane fragments of *A. actinomycetemcomitans* were prepared as described previously (1, 9, 10, 13). Briefly, 200 ml late logarithmic-phase cells were harvested, washed with phosphate buffered saline (PBS, 10 mM sodium phosphate, 150 mM sodium chloride, pH 7.4) and resuspended in 2.0 ml of 10 mM 4-(2-hydroxyethyl)-1-piperazineethanesulfonic acid (HEPES, pH 7.4) with 1 mM phenylmethylsulphonyl fluoride (PMSF, USB Corporation, Cleveland, OH) and Pierce protease inhibitors with EDTA (ThermoFisher Scientific, Rockford, IL). Bacteria were lysed in a French pressure mini-cell by three cycles of 9,000 p.s.i. (62,100 kPa) at 4°C. Whole cell lysates were centrifuged at 10,000 × *g* for 30 min to remove cell debris, followed by ultra-centrifugation at 100,000 × *g* for 1 h to separate the membrane and cytosol fractions. The membrane pellet was washed with HEPES, centrifuged and the resulting pellet was suspended in HEPES. The cytosolic fraction was re-centrifuged to removed contaminated membrane proteins. Protein concentrations were measured at 280 nm using an Evolution 201 UV-visible spectrophotometer (ThermoFisher Scientific, Waltham, MA).

### Biochemical analysis of EmaA

Equivalent amounts of protein were resuspended in buffer to a final concentration of 5% beta-mercaptoethanol, 0.06M Tris pH 6.8, 10% glycerol, 0.02% bromophenol blue, and 2% SDS; boiled for 5 min., and applied to wells of 4-15% polyacrylamide Tris-Glycine minigels (Bio-Rad, Hercules, CA) for electrophoresis (9, 10, 13). The separated proteins were transferred to 0.2 μm Immobilon-PSQ polyvinylidene difluoride (PVDF) membranes (MiliporeSigma, Burlington, MA) and probed with an anti-EmaA stalk monoclonal antibody (9). The immune complex was detected using HRP-conjugated goat anti-mouse IgG (Jackson Laboratory, Bar Harbor, Maine), and visualized using the SuperSignal West Pico plus chemiluminescent substrate (ThermoFisher Scientific, Waltham, MA).

### Transmission electron microscopy (TEM)

Bacteria were recovered from −80 °C and grown on TSBYE plates for 2-3 days, A single colony of each strain was inoculated into 10 ml of TSBYE broth and grown for 16h; the cultures were diluted 1:10 and grown for an additional 150 min. Bacteria were collected by centrifugation of 1 ml bacterial suspension at ×1,000 g for 1 min at 4°C and resuspended in 100 μl PBS pH7.4. Electron microscopy grids were prepared as previously described (4–6, 8). Briefly, a 5 μl aliquot of bacterial suspension was placed on either 300 or 200 mesh carbon-coated grids, and deep stained with NanoW™ (Nanoprobes, Yaphank, NY). For 3D electron tomography, the grids were pretreated with Poly-L-lysine (1000-5000 Da. Sigma, St. Louis, MO) and colloidal gold (SPI, West Chester, PA) to be used as fiducial markers.

### TEM Data acquisition

Data were collected using a Tecnai12 electron microscope (FEI, Hillsboro, OR) equipped with a LaB6 cathode (Kimball Physics, Wilton, NH), operated in point-mode (4, 21), a 2048 by 2048 pixel CCD camera with a pixel size of 14 μm, (TVIPS, Gauting, Germany) and a dual axis tilt tomography holder (Fischione, Export, PA). All images were recorded on the CCD camera at an acceleration voltage of 100 kV and a nominal magnification of 42,000, which corresponds to 0.308 nm pixel size on the specimen scale. Tomographic tilt series were acquired at least within a ±64° angular range in 2° angular intervals. Data were collected under low dose exposure conditions (10 e^-^/Å^2^ for 2D imaging and 3 e^-^/Å^2^ per image for the tomographic tilt series data) as previously described (8, 18).

### 3D reconstruction of the EmaA functional domain

Tomographic tilt series were processed using IMOD (22). Projections were initially aligned by cross correlation and further refined using fiducial markers located outside of the bacterial surface. EmaA functional domains were orientationally selected from the tomograms by marking two points along their long axis (11, 12, 17, 23) with the first point located at the tip of the adhesin. For each selected molecule, a tilt series of subprojections was extracted, subprojection angles were recalculated and subvolumes were reconstructed with the molecules’ long-axis oriented approximately parallel to the Y-axis using algorithms implemented in EMIRA (15). The 3D reconstruction algorithms based on Radon transforms, as implemented in EMIRA (15), contain an occupancy index that keeps track of the number of 2D transforms averaged into it and is used to determine the location of the missing data (24). Subvolumes were visualized in Chimera (25) and further aligned to a reference subvolume of the wild-type EmaA (6) using the “fit in map” command. Angles and shifts were applied to the subprojections and aligned subvolumes were reconstructed. The aligned subvolumes were grouped using Probabilistic Principle Component Analysis with Expectation Maximization (PPCA-EM), an algorithm to analyze 3D volumes with missing data, and Diday’s method of moving centers as clustering method implemented in EMIRA (15, 17, 24, 26, 27). Outliers (with sigma of 5) were identified and excluded from this stage of the analysis. A representative of each group was selected, and all representatives were aligned to each other. The members of each group were subsequently realigned to the aligned representative subvolume using the “fit in map” command in Chimera (25). Angles and shifts were applied to the subprojections and new aligned subvolumes were reconstructed. The newly aligned subvolumes were grouped anew using PPCA-EM followed by non-linear mapping and Diday’s method of moving centers as clustering method (15, 24, 26, 28). Final 3D structures were calculated by de novo reconstruction from 2D Radon transforms of the combined subprojections of all subvolumes in each group. In addition, to assess variability within groups, average subvolumes and variances were calculated from the reconstituted subvolumes after processing with PPCA-EM, which results in reconstituted subvolumes with the missing data filled in (15, 24). Averaged subvolumes and variances were low pass filtered and visualized in Chimera (25). Segmentations were performed in Chimera using initially a 2-step smoothing and finally a 3-step smoothing (29).

### Collagen binding assay

A polyclonal antibody was developed against a serotype a strain of *A. actinomycetemcomitans* as described before (3). The collagen binding activity of different strains was evaluated using 96-well ELISA plates, as described before (30). Briefly, Type V collagen from human placenta (Sigma Type IX, C3657, Sigma-Aldrich, St Louis, MO) was pre-solubilized in aqueous acid and diluted in carbonate coating buffer, pH 9.6. A 100 μl aliquot at 10 μg/ml was loaded per well, and stabilized overnight at 4°C. The wells were rinsed with PBS, pH 7.4 and blocked in PBS complemented with 0.5% bovine serum albumin. Approximately 1.0E7 bacteria per well were incubated for 1 h, washed with PBS, and incubated with a suspension of 4.6 μg/ml anti-serotype a polyclonal antibody. After 1 h incubation, plates were washed with PBS containing 0.05% Tween 20 and incubated for 1 h with the secondary antibody, horseradish peroxidase-conjugated (HRP) goat anti-rabbit immunoglobulin (Jackson Laboratory, Bar Harbor, ME). Immunoglobulin complexes were detected using citrate-phosphate buffer pH 5.0 containing 0.04% O-phenylenediamine and 0.012% hydrogen peroxide. The reaction was stopped by addition of 50 μl of 4 M H_2_SO_4_ and, absorbance was measured at 490 nm. Data were analyzed using paired *t*-test with GraphPad Prism 7.0a (GraphPad, San Diego, CA), and *P* < 0.05 was considered significant.

### Biofilm formation assay

Biofilm formation was evaluated using 96-well cell culture plates with Nunclon Delta cell culture treatment (ThermoFisher Scientific) as described before (3). Briefly, bacteria were recovered from −80 °C and grown on TSBYE plates for 3 days. A single colony of each strain was inoculated into 10 ml of TSBYE broth and grown overnight; subsequently, the cultures were diluted 1:10 and grown for one doubling time (∼ 150 min). Each bacterial strain was load with ∼1.0E6 bacteria in 200 μl/per well, and grown for either 24 h, 48 h and 7 2h. The top 100 μl culture medium was gently replaced with fresh medium every 24 h. Wells were washed with PBS pH 7.4 three times, stained with 0.35% crystal violet in 19% ethanol and 1% methanol, washed to remove extracellular dye, and de-stained using 95% ethanol. Semi-quantification was performed using a spectrophotometer (absorbance at 562 nm). Proper dilution of the crystal violet was required to avoid instrument saturation. Data were analyzed using a paired *t*-test with GraphPad Prism 9.3.1 (GraphPad, San Diego, CA), and *P* < 0.05 was considered significant.

## ACKNOWLEDGMENTS

We appreciate the technical support and discussions from David Danforth, Marcella Melloni, Jake Tristano, Claire Brooks and Alison Watson.

## FUNDING INFORMATION

This work was supported, by National Institute of Health/National Institute of Dental and Craniofacial Research (Grant DE024554 funded to Teresa Ruiz & Keith P. Mintz), as well as NIH/National Institute of General Medical Sciences (Grant GM078202 funded to Michael Radermacher), and National Science Foundation (Grant DBI 1660908 funded to Michael Radermacher).

## SUPPLEMENTAL MATERIALS

1. Movie1.mp4: surface representation of subvolume averages of the functional N-terminus of the b-EmaA expressed in the transformed serotype a strain. EmaA subvolumes reconstructed tomograms (yellow surface) versus the wild type (WT) b-EmaA (black mesh).
2. Movie2.mp4: segmentation of subvolume averages of the functional N-terminus of the b-EmaA expressed in the transformed serotype a strain. WT represents the wild type b-EmaA.

